# MICU1 occludes MCU in the mitochondrial calcium uniporter complex

**DOI:** 10.1101/2021.11.15.468501

**Authors:** Chen-Wei Tsai, Anna Van Keuren, John Bankston, Zhiwei Ma, Ming-Feng Tsai

**Affiliations:** Department of Physiology and Biophysics, University of Colorado Anschutz Medical Campus, Aurora, CO 80045; Dalton Cardiovascular Research Center, University of Missouri, Columbia, MO 65211

**Author notes:** Correspondence should be addressed to Ming-Feng Tsai.

## Abstract

The mitochondrial calcium uniporter imports cytoplasmic Ca^2+^ into the mitochondrial matrix to regulate cell bioenergetics, Ca^2+^ signaling, and apoptosis. The uniporter contains the pore-forming MCU subunit, an EMRE protein that binds to MCU, and the regulatory MICU1/MICU2 subunits. Structural and biochemical studies have suggested that MICU1 gates MCU by blocking and unblocking the Ca^2+^ pore. However, mitoplast patch-clamp experiments argue that MICU1 does not block Ca^2+^ transport but instead potentiates MCU. To address this direct clash of proposed MICU1 function, we applied purified MICU1 to Ca^2+^-conducting MCU-EMRE subcomplexes in outside-out patches excised from *Xenopus* oocytes. MICU1 strongly inhibits Ca^2+^ currents, and the inhibition is abolished by mutating an MCU-interacting K126 residue in MICU1. Further experiments show that MICU1 block was not observed in mitoplasts because MICU1 dissociates from the uniporter complex. These results firmly establish that MICU1 shuts the uniporter in resting cellular conditions.

## Introduction

The mitochondrial Ca^2+^ uniporter is a multi-subunit Ca^2+^ channel complex in the inner mitochondrial membrane (IMM). Its transmembrane (TM) region contains the MCU subunit, which forms a tetrameric Ca^2+^ pore^1–3^, and the EMRE protein that binds to MCU to stabilize the pore’s conducting conformation^4,5^. The MICU subunits, which include MICU1, MICU2, and the neuron-specific MICU3 (not discussed here), contain two canonical EF-hands and dwell in the intermembrane space (IMS) to regulate uniporter activity^1–3,6^. MICU1 regulation of the uniporter is important for normal physiology, as MICU1 mutations have been linked to severe neuromuscular disorders in humans^7,8^. The exact regulatory function of MICU1, however, is currently under debate.

Biochemical and functional analyses^1–3,9,10^ have suggested a mechanism that MICU1 (but not MICU2) binds to the Ca^2+^-coordinating Asp ring (D261 in human MCU) at the cytoplasmic entrance of the MCU pore to block Ca^2+^ flux at resting levels of cytoplasmic Ca^2+^ (100 – 200 nM). Upon elevation of local Ca^2+^, MICU1 dissociates from the pore to enable Ca^2+^ permeation (Fig. 1A). This “occlusion model” is supported by multiple key results: first, co-immunoprecipitation shows that human MCU (hMCU) can complex with MICU1 in the absence of EMRE, and that such MCU-MICU1 interaction is disrupted by a D261A mutation in hMCU^9^; second, a quantitative ^45^Ca^2+^ flux assay shows that substituting wild-type (WT) hMCU with the D261A mutant in WT HEK293 cells leads to robust mitochondrial Ca^2+^ uptake at resting Ca^2+^, phenocopying MICU1-knock out (KO)^9^; and third, MICU1 suppresses inhibition of MCU by Ru360, which binds to D261^10^. Recent cryo-electron microscope (cryo-EM) studies appear to provide the structural basis of the occlusion model^1–3^. In low Ca^2+^, MICU1 uses a 5-residue Arg/Lys ring to surround MCU’s D261-ring to block the channel (Fig. 1B). Ca^2+^ binding to MICU1 induces conformational changes that disrupt MICU1-MCU interactions to open the pore (Fig. 1B). Functional work^3^ further shows that mutations in the Arg/Lys ring abolish MICU1 block of uniporter Ca^2+^ transport in resting cellular Ca^2+^.

**Figure 1.**
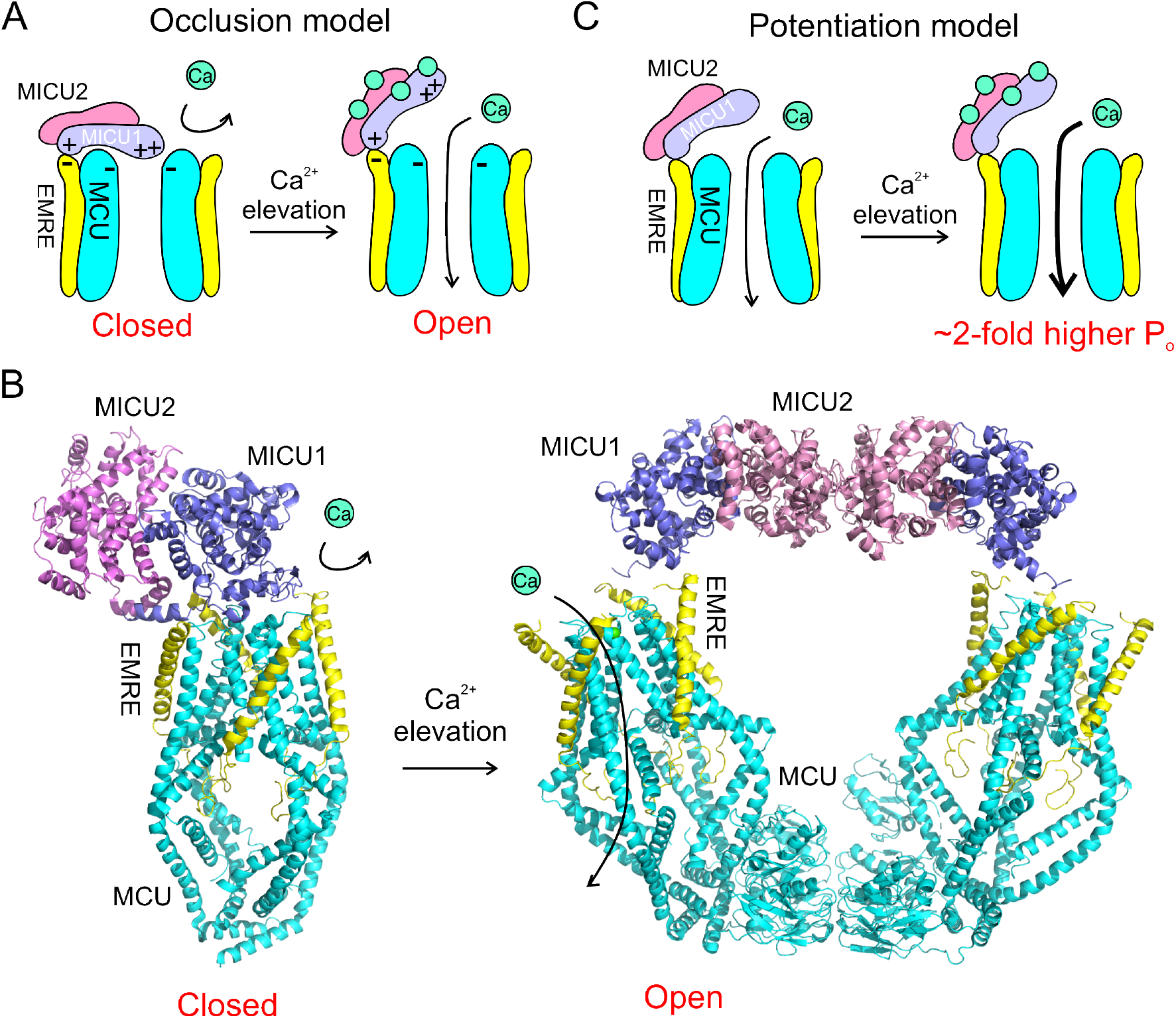
Two models for Ca^2+^-dependent gating of the uniporter. **(A)** The occlusion model. The cartoon illustrates that MICU1 interacts with both EMRE and MCU via electrostatic interactions in low Ca^2+^. In high Ca^2+^, the MCU-MICU1 interaction is disrupted so that Ca^2+^ can pass the pore, but the EMRE-MICU1 interaction remains intact to keep MICU1 associated within the uniporter complex. **(B)** Uniporter holocomplex structures in low (PDB: 6WDN)/high Ca^2+^ (6WDO). These structures recapitulate the key propositions in the occlusion model that MICU1 blocks MCU in low Ca^2+^ (left) but swings away to open the pore in high Ca^2+^ (right). **(C)** The potentiation model. This model proposes that MICU1 does not block MCU, but upon Ca^2+^ binding, MICUs increase the open probability of MCU by 2-fold via EMRE-dependent allosteric mechanisms.

The occlusion model, however, directly clashes with results in a landmark study, which employs patchclamp of mitoplast (a submitochondrial vesicle rid of the outer membrane) to demonstrate that the uniporter is a *bona fide* Ca^2+^ channel^11^. In this work, it was shown that the uniporter can conduct Na^+^ in 0 Ca^2+^, apparently contradicting the idea that MICU1 physically occludes the MCU pore under such a 0 Ca^2+^ condition. Expanding on these observations, a recent electrophysiological work^12^ further proposes a “potentiation model” for MICU1 (Fig. 1C). Based on the finding that MICU1-KO reduces uniporter Ca^2+^ currents by ~50% in mitoplasts, it was argued that MICU1’s function is to potentiate the uniporter rather than to block MCU. The goal of this work is to understand why rigorous work from multiple labs over the last two decades produce two entirely opposite models underlying Ca^2+^-dependent gating of the uniporter by MICU1.

## Results

### Purified, EF-hand mutated MICU1 blocks MCU Ca^2+^ currents

The critical difference between the occlusion and potentiation models is that the former demands the presence of a MICU1-mediated, blocked state, while the latter argues that MICU1-block is nonexistent. Thus, the key to determine which of these two models is correct is to examine if there is a blocked state. We thus test if purified MICU1 can inhibit MCU currents, analogous to classical K^+^-channel work^13,14^ showing that purified neurotoxins (*e.g*. charybdotoxin) or synthesized ball peptides (as in the ball-and-chain mechanism) block heterologously expressed K^+^ channels. Such a strategy was deemed feasible as cryo-EM work has already shown that purified MICU1 can bind to the MCU-EMRE pore subcomplex reconstituted in lipid bilayers^1^. We excised outside-out patches from *Xenopus* oocytes expressing a human MCU-EMRE (hME) fusion protein in the cellular membrane. Our previous work^15^ demonstrated that hME exhibits similar electrical properties as the uniporter recorded in mitoplast patch-clamp^11^. At negative voltages, hME conducts inward Ca^2+^ currents that are strongly and reversibly inhibited by Ru360 (Fig. 2A). We then perfused MICU1_EF_, which includes 4 mutations (D231A, E242K, D421A, E432K) to eliminate Ca^2+^ binding to the EF hands to mimic low Ca^2+^ conditions. It has been shown qualitatively that transiently expressed MICU1_EF_ in HEK cells suppresses mitochondrial Ca^2+^ uptake^16^, a result recapitulated here using quantitative ^45^Ca^2+^ flux (Fig. 2B). Strikingly, applying MICU1_EF_ reduces macroscopic hME Ca^2+^ currents by 86 ± 7%, with a current decay time constant of 44 ± 12 s (i-ii, Fig. 2A). Moreover, in recordings wherein single-channel transitions can be observed (iii-iv, Fig. 2A), MICU1_EF_ greatly reduces hME open time and open probability. The inhibition is not reversible with a 3 – 5 min wash. These results are fully explained by a molecular picture that purified MICU1_EF_ binds to hME with a slow on-rate to block hME in a highly stable MICU1_EF_-hME complex (scheme I).

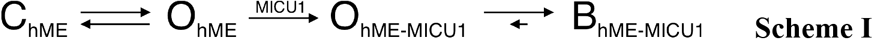

**Figure 2.**
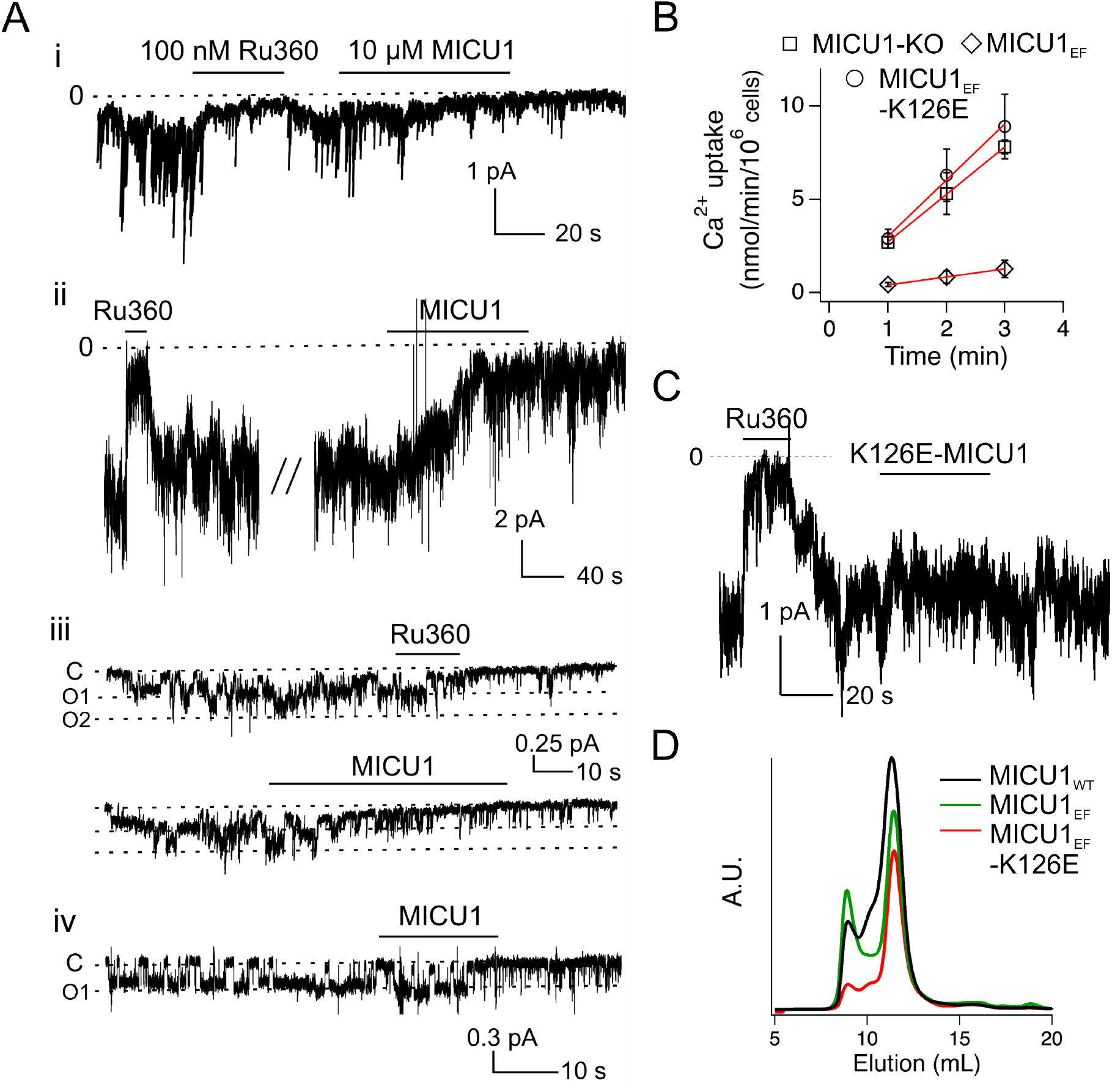
The effect of MICU1_EF_ on uniporter Ca^2+^ currents. **(A)** The effects of perfusing MICU1_EF_ to patches containing Ca^2+^-conducting hME. Membrane potentials were clamped at −60 to −100 mV. MICU1_EF_ was tagged with an N-terminal maltose binding protein to facilitate proper protein folding in the periplasmic space of *E. coli*. In all recordings, [MICU1] and [Ru360] are 10 μM and 100 nM, respectively. N = 7 (4 macroscopic recordings and 3 recordings with 1 – 2 channels). **(B)** Mitochondrial uptake of ^45^Ca^2+^ in MICU1-KO cells. The experiments were performed in the presence of 250 nM Ca^2+^, with MICU1-KO transiently expressing indicated MICU1 mutants. N = 5 **(C)** Mutational effects of K126E on MICU1 function. Voltage was clamped at −80 mV. N = 3. **(D)** Size exclusion profiles of WT MICU1 and MICU1 mutants. N = 4 – 8.

To test if MICU1_EF_ inhibits hME via the blockade interactions observed in uniporter-complex structures (Fig. 1B), we mutated MICU1’s K126 residue, which contacts MCU’s D261 ring to block the pore. ^45^Ca^2+^ flux shows that K126E abolishes the ability of transiently expressed MICU1_EF_ to inhibit mitochondrial Ca^2+^ uptake in HEK cells (Fig. 2B). Moreover, perfusing K126E-MICU1_EF_ leaves macroscopic hME Ca^2+^ currents unchanged, indicating that K126E indeed abolishes MICU1_EF_ block (Fig. 2C). Size-exclusion chromatography shows that purified K126E-MICU1_EF_ preserves the proper biochemical behavior of MICU1_EF_ (Fig. 2D), suggesting that K126E does not perturb MICU1 folding. Taken together, our electrical recordings demonstrate that MICU1_EF_ directly inhibits MCU via a K126-mediated interaction observed in the blocked state of the uniporter-complex structure in 0 Ca^2+^ (Fig. 1A-B). These results thus present novel electrophysiological evidences that support the occlusion mechanism and cannot be explained by the potentiation model.

### No MICU1 block of MCU in mitoplasts because MICU1 is dissociated

We then sought to understand why MICU1 block was not observed in mitoplasts^11^. As MICU1 contacts the uniporter’s TM subunits via electrostatic interactions^3,9^, it is possible that high-salt treatments in the mitoplast production procedure might lead to MICU1 dissociation from the TM subunits^11^. Accordingly, we employed quantitative Western blots to compare the MICU1/MCU signal ratio in mitochondria vs mitoplasts produced following published procedures in the original mitoplast patch-clamp experiments^11^. Results show a 30% reduction of MICU1/MCU ratio in mitoplasts (Fig. 3A), suggesting that ~30% of MICU1 dissociates into the bulk solution during mitoplast formation.

**Figure 3.**
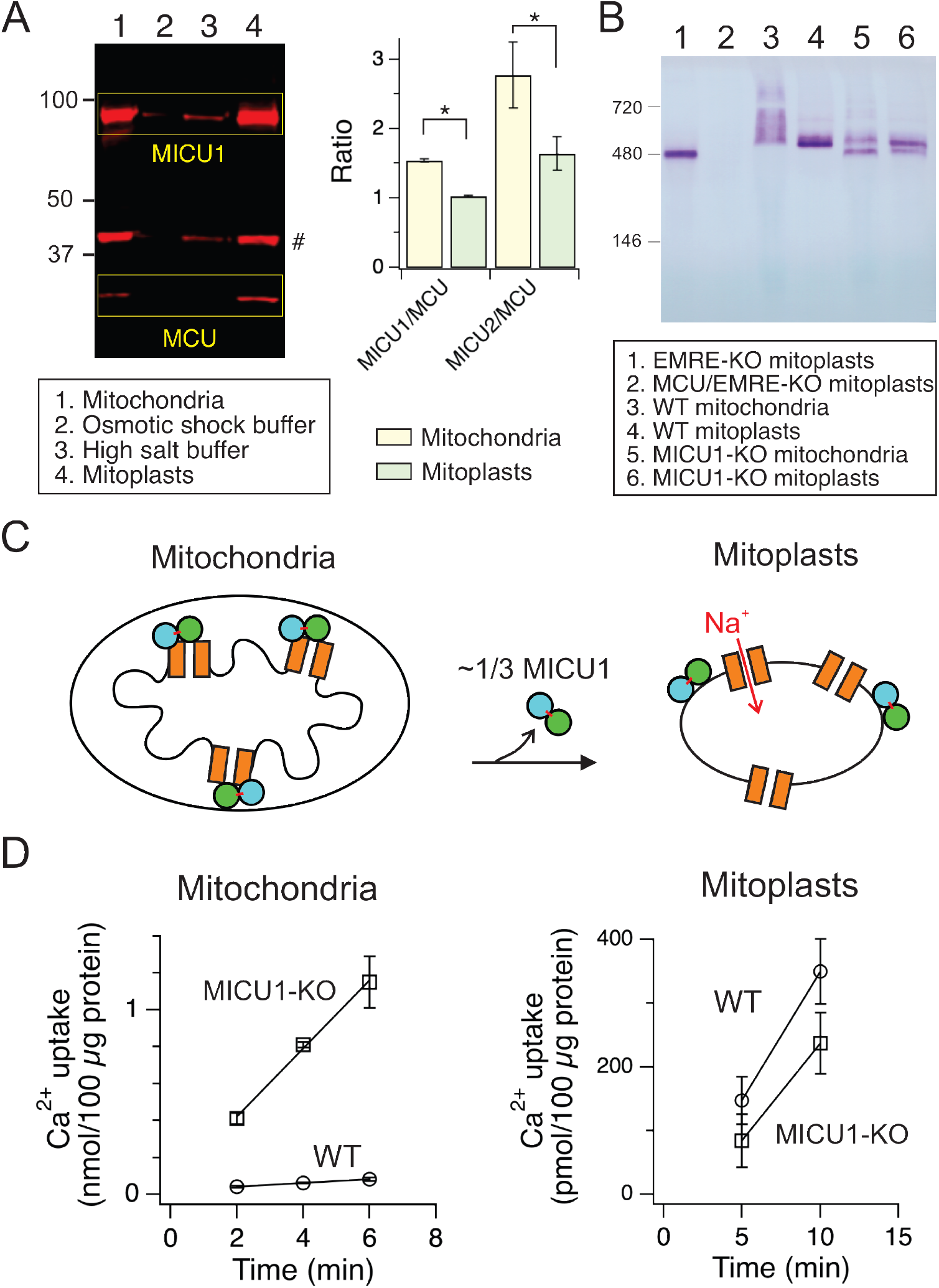
Biochemical and functional characterization of the uniporter in mitoplasts. **(A)** Changes of relative MICU/MCU levels during mitoplast preparation. The *#* sign marks a non-specific band. N = 4 – 6. **(B)** BN-PAGE analysis of the uniporter complex. N = 4. **(C)** A cartoon explaining why MICU1 block is not observed in mitoplasts. The connected green-cyan balls represent the MICU1-MICU2 heterodimer connected by an intersubunit disulfide. **(D)** Deregulation of the uniporter in mitoplasts. The experiments were performed in the presence or absence of Ru360 to allow the calculation of uniporter-mediated, Ru360-sensitive Ca^2+^ uptake in mitochondria or mitoplasts. N = 4 – 6.

Since the uniporter preferentially dwells in areas near the inner/outer membrane contact sites^3^, the disruption of such contact sites in mitoplasts could effectively dilute the uniporter in a much bigger surface area, rendering it more difficult for dissociated MICU1 to bind back to the TM region. We thus wondered if a portion of remaining MICU1 in mitoplasts might only attach to the IMM without being associated with the uniporter (MICU1 is a peripheral membrane protein). To this end, the uniporter complex was analyzed using blue native PAGE (Fig. 3B). The uniporter in WT mitochondria runs as a smear (lane 3), likely due to the presence of uniporter complexes binding different numbers of MICU1. Accordingly, the smear is eliminated by MICU1-KO (lanes 5–6). MICU1-KO reduces EMRE expression^17^, thus creating EMRE-bound (upped band, lanes 5–6) and EMRE-free (lower band) complexes. The latter is also observed in EMKE-KO samples (lane 1). In WT mitoplasts, the uniporter runs as a single band without a smear (lane 4), and the band position is consistent with the EMRE-bound, MICU1-free uniporter (lane 5–6). These results thus argue strongly that the mitoplast production process leads to dissociation of MICU1 from the uniporter complex (Fig. 3C), creating a subpopulation of MICU1-free uniports that can conduct Na^+^ currents in 0 Ca^2+^.

To further test the scheme in Fig. 3C using functional assays, we analyzed ^45^Ca^2+^ flux into mitochondria or mitoplasts. In 250 nM Ca^2+^, mitochondria from MICU1-KO HEK cells take up much more Ca^2+^ than WT mitochondria (Fig. 3D), as expected from the loss of MICU1 block in the occlusion model (Fig. 1A). However, WT mitoplasts take up comparable amount of Ca^2+^ in 250 nM Ca^2+^ as MICU1-free mitoplasts (Fig. 3D), thus demonstrating that MICU1 regulation of the uniporter is severely perturbed in mitoplasts.

## Discussion

The uniporter complex emerges in eukaryotic evolution with only MCU and MICU1 subunits^18^, suggesting that MICU1 must regulate MCU through direct physical contacts. Indeed, the MCU-MICU1 interaction is so conserved that human MICU1 can complex with MCU homologues in *A. thaliana* (plant), *D. discoideum* (amoeba)^9^, or *T. castaneum* (beetle)^2^. When MCU structures were first determined, its small and flat IMS surface makes it rather obvious that docking of MICU1 onto this surface would block ion permeation through the Ca^2+^ pore^19^. This is indeed confirmed by subsequent biochemical and structural work summarized in the Introduction, leading to the establishment of the occlusion mechanism (Fig. 1A). In the potentiation model, perhaps realizing that MICU1 would unlikely increase uniporter activity by binding to MCU’s IMS surface, it was proposed that MICU1 potentiates MCU allosterically through EMRE (Fig. 1C)^12^. Such an argument, however, lacks apparent structural basis, as EMRE-MCU interactions and the MCU pore conformation in the MCU-EMRE-MICUs structure appear identical in low and high Ca^2+^ conditions^1–3^. Moreover, EMRE is only present in animals — an EMRE-dependent potentiation mechanism would imply that the highly conserved MICU1 protein, which co-evolves with MCU since early eukaryotes^18^, actually has no function in the uniporter in non-metazoan organisms.

Our electrical recordings, coupled with structural information, directly demonstrate MICU1 block of the MCU pore in real time, a phenomenon that the potentiation model cannot explain. Moreover, our biochemical and functional investigations resolve the longstanding puzzle about why the uniporter can conduct Na^+^ without Ca^2+^ in mitoplast membranes. However, we note that it remains unclear why MICU1-KO reduces uniporter Ca^2+^ currents in mitoplast patch-clamp experiments^12^. Our work (under consideration elsewhere) shows that MICU1-KO indeed reduces uniporter-mediated ^45^Ca^2+^ uptake, but such reduction is caused by EMRE downregulation (rather than loss of MICU1 potentiation), because transient EMRE expression in MICU1-KO HEK cells restores uniporter activity to WT-cell levels. However, mitoplast patch-clamp shows that overexpressing EMRE does not further enhance macroscopic Ca^2+^ currents in MICU1-free mitoplasts^12^. We do not know how to explain the discrepancy, but would like to point out that proper expression of EMRE in MICU1-KO cells must be carefully confirmed with quantitative methods, given EMRE expression is strongly suppressed in these cells.

## Materials and Methods

### Electrophysiology

*Xenopus* oocytes were injected with 60 ng of cRNA encoding the human MCU-EMRE fusion protein (hME) and were incubated in an ND96 solution (2 mM KCl, 96 mM NaCl, 1.8 mM CaCl_2_, 1 mM MgCl_2_, 5 mM HEPES, pH 7.4-NaOH) at 17 °C as described in our previous work^15^. Patch-clamp recordings were performed 3 – 5 days after cRNA injection. Borosilicate pipettes were produced using a Sutter P1000 pipette puller, and were fire polished with a homemade microforge, leading to a pipette resistance of 3 – 5 MΩ. In all experiments, the pipet solution contains 90 mM sodium gluconate, 10 mM NaCl, 5 mM EGTA, 20 HEPES, pH 7.6-NaOH. Outside-out patches were excised from oocytes in a continuous perfusion ND96 bath, and then moved to the outlet of a fast solution change system (Warner SF-77B) to be perfused with a Ca-100 solution (100 mM CaCl_2_, 20 mM HEPES, pH 7.6-NaOH). Currents of hME were recorded at room temperature with a HEKA EPC-10 amplifier, filtered at 100 Hz with an eight-pole Bessel filter (Warner), and digitized at a frequency of 500 Hz. The membrane potential was clamped at −60 to −100 mV (more negative voltages were used when the seal was more stable). Negative (inward) currents are defined as currents from the bath to the cytoplasmic side of outside-out membranes. 100 nM Ru360 or 10 μM MICU1 were diluted in Ca-100 and was applied to the patches via the SF-77B fast solution change system.

### Protein purification

DNA encoding N-terminally maltose binding protein (MBP)-tagged MICU1 proteins was cloned into a pET21b vector. Transformed *E. coli* BL21 cells were inoculated into terrific broth, grown to an OD of 1.0 at 37 °C, and then induced for protein expression by adding 0.1 mM IPTG for 2 h at 37 °C. Cells were pelleted, resuspended in a breaking buffer (BB, 500 mM NaCl, 25 mM HEPES, pH 7.5-NaOH) supplemented with 1 mg/mL leupeptin/pepstatin and 1 mM PMSF. After lysis by sonication, the lysate is clarified by centrifugation at 16,000 g for 30 min at 4 °C. The lysate wan then loaded onto a cobalt-affinity column. The column was washed and eluted with 10 mM and 200 mM imidazole in BB, respectively. The protein was then concentrated and further purified via a Superdex 200 Increase size-exclusion column equilibrated with BB. Elution volume 10.5 – 12 mL was collected and concentrated down to 150 μM. The concentrated stock was stored at −80 °C and diluted to Ca-100 right before electrophysiological experiments to be used within 4 hr. The freeze-thaw process and leaving protein in Ca-100 for 4 hr in room temp does not affect the size-exclusion profiles. The yield of WT-MICU1, MICU1_EF_, and K126E-MICU1_EF_ are 15 mg, 5 mg, 2.5 mg/L *E.coli*.

### Cell culture and molecular biology

HEK 293T cells were cultured in DMEM with 10% FBS in a 5% CO_2_ incubator at 37 °C. MICU1-KO was achieved by CRISPR-Cas9, and has been validated in our previous publication^17^. The genes encoding MICU1 mutants were cloned into a pcDNA3.1(+) vector for transient expression using Lipofectamine 3000 (Life Technologies). Site-directed mutagenesis was performed using a Quick Change kit (Agilent) with the mutations verified by Sanger sequencing.

### Mitochondria/mitoplast production, Western blot, and blue-native (BN)-PAGE

Mitochondria and mitoplasts were produced following the protocols described in the original mitoplast patch-clamp paper^11^ with some minor modifications. Briefly, HEK cells suspended in a mitochondrial resuspension buffer (250 mM sucrose, 5 mM HEPES, 1 mM EGTA, pH 7.2-KOH) were disrupted by passing through 27.5 gauge needles 10 – 20 times. Mitochondria were pelleted by differential centrifugation (1000 g once followed by 13,000 g 3 times) at 4 °C. To obtain mitoplasts, mitochondria were resuspended in an osmotic shock buffer (5 mM sucrose, 5 mM HEPES, 1 mM EGTA, pH 7.2-KOH) for 10 min on ice, spun down at 13,000 g for 10 min at 4 °C, resuspended in a high-salt buffer (750 mM KCl, 100 mM HEPES, 1 mM EGTA, pH 7.2-KOH) for 30 min on ice, and then pelleted by centrifugation at 13,000 g for 10 min at 4 °C.

To perform Western blots as in Fig. 3A, samples obtained in the mitoplast production process were subjected to SDS-PAGE and transfer to PVDF membranes. The membranes were treated with both anti-MICU1 (Sigma, HPA034780, 1:5,000) and anti-MCU (Cell Signaling, D2Z3B, 1:10,000) antibodies at 4 °C for overnight, followed by a IRDye 680 goat anti-rabbit secondary antibody (Li-Cor, 1:10,000) in room temperature for 1 h. The image was acquired using a Li-Cor Odyssey CLx imager.

BN-PAGE was performed using the Novex NativePAGE Bis-Tris gel system (Life Technologies). 200 μg of mitochondria or 45 μg of mitoplasts was dissolved in 20 μL of NativePAGE buffer supplied with 4% digitonin, 4% glycerol, and 0.5% G-250. Samples were loaded onto 4 – 16% Bis-Tris gels, subjected to electrophoresis at 4 °C, and transferred to PVDF membranes. The membranes were de-stained by a solution containing 25% methanol and 10% acetic acid, blocked with 5% milk in TBS (140 mM NaCl, 3 mM KCl, 25 mM HEPES, pH 7.4-HCl), and then incubated with anti-MCU antibody (Cell Signaling, D2Z3B, 1:5000) in TBST (TBS + 0.05% Tween 20) for overnight at 4 °C, followed by anti-rabbit alkaline phosphatase-conjugated secondary antibody (Promega, S3731, 1:5,000 in TBST) at room temperature for 1 h. Colorimetric detection was performed by adding NBT/BCIP substrates (Life Technologies) to the membranes.

### Mitochondrial and mitoplast ^45^Ca^2+^ uptake assays

To measure ^45^Ca^2+^ flux into mitochondria, mitochondria were isolated using procedures described above and quantified using the BCA assay (Pierce). 350 μg of mitochondria were resuspended in 1 mL of a wash buffer (WB, 120 mM KCl, 25 mM HEPES, 2 mM KH_2_PO_4_, 1 mM MgCl_2_, pH 7.2-KOH), before being pelleted. ^45^Ca^2+^ uptake was initiated by resuspending mitochondria in 350 μL of a low Ca^2+^ recording buffer (LCRB, 120 mM KCl, 25 mM HEPES, 2 mM KH_2_PO_4_, 5 mM succinate, 1 mM MgCl_2_, 0.69 mM EGTA, 0.5 mM CaCl_2_, pH 7.2-KOH), and was terminated at 2, 4, and 6 min by adding 100 μL of the suspension into 5 mL of ice-cold WB, and filtered through nitrocellulose membranes. The membranes were extracted in a scintillation cocktail for scintillation counting. Th rate of mitochondrial Ca^2+^ uptake was obtained by linear fit of data points, with detailed calculation methods provided in our previous work^9^. Mitoplast ^45^Ca^2+^ flux was performed in a similar manner, with the following difference: Ca^2+^ uptake was quenched by adding 100 μL mitoplasts to 1 mL of ice-cold WB supplemented with 200 nM Ru360 and 5 μM CGP-37157, and external ^45^Ca^2+^ was removed by spinning down mitoplasts at 13,000 g for 10 min at 4 °C. All ^45^Ca^2+^ uptake experiments were performed in the presence or absence of 200 nM Ru360 to allow isolation of uniporter-specific mitochondrial Ca^2+^ uptake.

### Data analysis and statistics

Data points were presented as mean ± S.E.M. Statistical analysis was performed using t-test, with significance defined as *p* < 0.05. All experiments were done in at least 3 independent repeats. Analysis of electrophysiological data was done using Igor Pro 8 (WaveMetrics). Western blot quantification was done using the Li-Cor Image Studio software (version 5.2).

## Acknowledgements

We thank Madison Rodriguez and Dr. Han-I Yeh for technical assistance, and Dr. Christopher Miller for critical reading of the manuscript. We thank Dr. Yuriy Kirichok and Dr. Vivek Garg for offering two rounds of in-depth reviews and multiple days of intense but respectful email debates, leading to strongly improved manuscript. We thank Dr. György Hajnóczky for discussions during the development of this work. CT, AV, and MF are supported by the NIH grant R01-GM129345.

## Author Contributions

MT conceptualized the project; CT, AV, and ZM performed research. CT and AV analyzed data. JB supplied key resource. MT wrote the paper.

